# Target Distance from the Visual Field and Increased Age Affect Visual Search Efficiency: Behavioral and Modeling Evidence

**DOI:** 10.1101/2024.03.28.587192

**Authors:** Fatemeh Akbari, Samaneh Asivandzadehchaharmahali, Alireza Tanha, Abdolvahed Narmashiri

## Abstract

Previous research has demonstrated that visual search is influenced by environmental factors, but the effects of specific variables, such as target distance from the visual field center and age, are not well understood. To address this issue, we aim to investigate their impact on visual search task. Participants engaged in target-present and target-absent trials, revealing distinct patterns in search times. Behavioral data and drift-diffusion modeling (DDM) showed that increasing the target’s distance from the center of the visual field significantly reduces search efficiency. Additionally, age negatively impacts search performance, with older individuals exhibiting reduced efficiency. This comprehensive examination contributes to understanding cognitive mechanisms in visual processing. These findings highlight the importance of considering spatial and age-related factors in visual search tasks.

## Introduction

Humans often encounter the challenge of rapidly identifying valuable objects among numerous irrelevant ones in their surroundings during conjunction visual search, wherein individuals identify a target stimulus among distractors based on multiple features ^1^. Although conjunction visual search can be elucidated through the serial processing of features like color, size, or orientation across the visual field, the precise roles of several critical factors in this type of search remain uncertain.

The efficiency of visual search is influenced by factors such as the presence or absence of a target, the complexity of the search display ^2^, target distance from the central visual field ^3–5^, and individual differences, including age-related cognitive dynamics ^6,7^. One crucial aspect of conjunction visual search is the time it takes to detect a target, particularly when the target is present compared to when it is absent. Previous research has highlighted the role of attention and cognitive processes in shaping search times ^8,9^. By exploring the temporal aspects of search tasks, we aim to contribute to a deeper understanding of the underlying mechanisms that govern visual search behavior. Furthermore, investigating the precision of target localization adds another layer of complexity to our exploration. Understanding how accurately individuals can pinpoint the location of a target in a visual field provides insights into the spatial dynamics of visual search ^10–12^. This aspect is particularly relevant in scenarios where multiple features need to be processed conjointly to identify the target, forming the basis of conjunction visual search. Age-related differences in cognitive processing have been well-documented, and visual search is no exception. Previous studies suggest that aging can impact various aspects of visual attention, including search efficiency and the ability to filter out irrelevant information ^7,13,14^. It remains unclear whether the location of the target in the visual field and age-related dynamics impact search time in finding the target during conjunction visual search.

To address this issue, we conducted a behavioral experiment using conjunction visual Search. This task comprised two types of trials: those with a target presented in the array (target-present trials) and those without a target in the array (target-absent trials). Firstly, we examined the effect of trial type on search time. Secondly, we evaluated the precision of target localization and elucidated age-related dynamics on search time during conjunction visual search. Briefly, the results indicate that longer search times occurred in target-absent trials with larger set sizes, while accuracy was maintained. Conversely, larger set sizes in target-present trials resulted in reduced accuracy, indicating a higher cognitive load. The study also observed longer search times with greater distances and set sizes from the central visual field, underscoring the complex interaction between spatial factors and set size in target localization. Additionally, older age was correlated with longer search times in target-absent trials, suggesting difficulties in confirming absence.

## 1. Method

### 2.1. Participants

All participants reported normal or corrected-to-normal vision and were screened for mental illnesses, personality issues, drug or alcohol abuse, neurological disorders, and epilepsy. This study is supported by the School of Cognitive Science at the Institute for Research in Fundamental Sciences (IPM). Our research strictly followed the ethical guidelines outlined in the Declaration of Helsinki. Before participating in the study, all participants provided written informed consent. The Experiment had 46 participants (6 men, 40 women, age mean: 39/55). The gender of participants was determined based on self-report.

## Stimuli

The visual search task employed a paradigm designed to investigate distinctions between search processes ^15,16^. Commencing each task involved fixing the gaze on a central reference point displayed on the screen. The trial’s target was indicated by briefly superimposing the fixation cross with the target itself, succeeded by the presentation of an array of stimuli incorporating both the target and distractors (Fig. 1A). The task featured a conjunction-style search array, wherein the target shared a common feature property with the distractors, either in terms of shape (’T’ or ’L’) or color (red or green) which appeared in one of the 81 locations on the screen (Supplementary Fig. 1A). The search arrays varied in size, comprising a total of 6, 12, 24, or 48 stimuli. Participants were tasked with acquiring and maintaining fixation cross on the target until the spacebar was pressed. Their objective was to signal the presence or absence of the target within the array by pressing one of two buttons. Notably, the target was present in half of the trials (Fig. 1A).

**Fig. 1.**
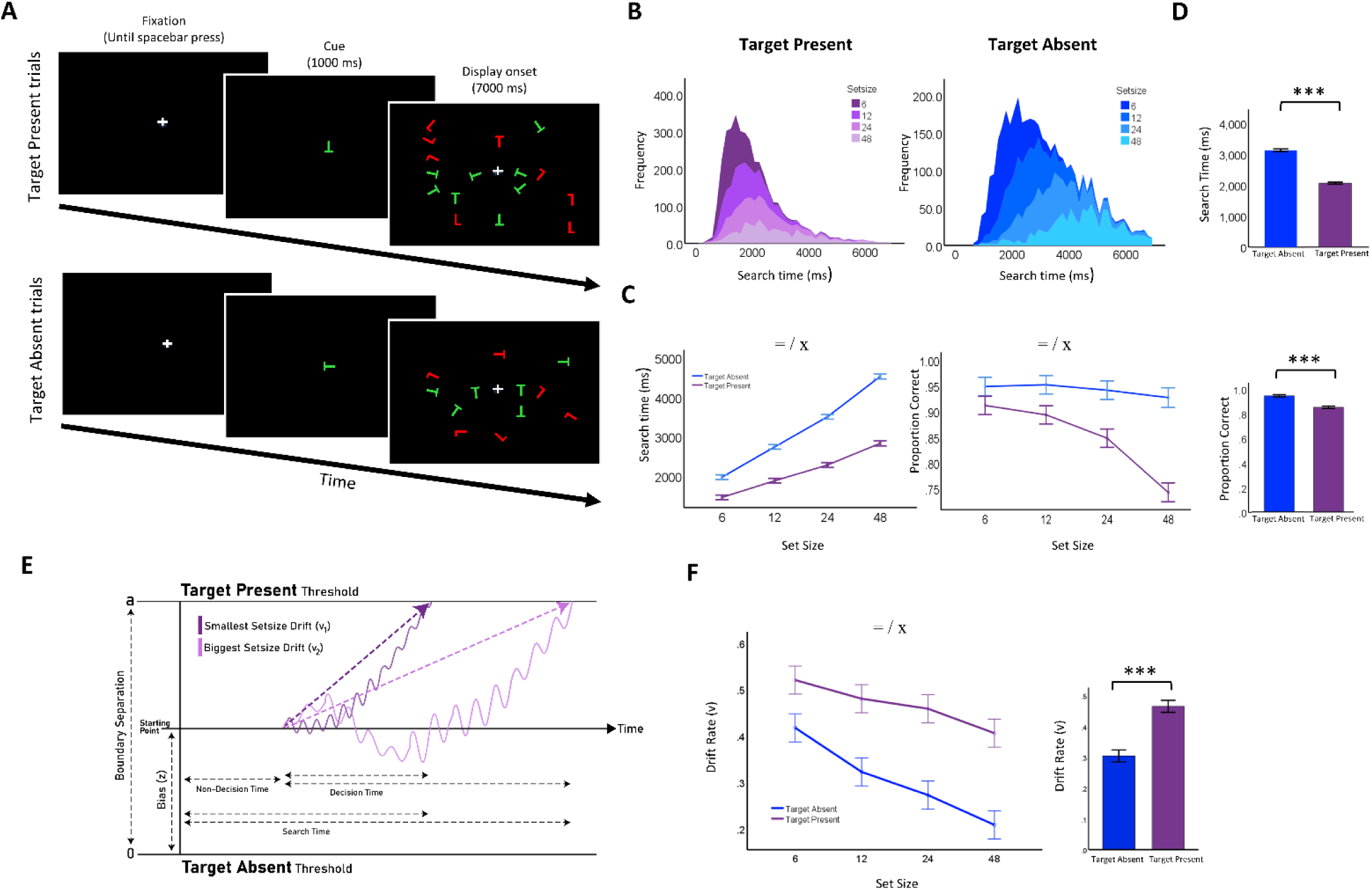
Visual search paradigm, task design, behavioral, and modeling results. A) The search task comprised target-present (TP) and target-absent (TA) trials. In each trial, a variable number of objects (6, 12, 24, or 48) were presented on the screen. In TP trials, only one of the displayed objects served as the target, while in TA trials, there was no target. Subjects were required to press two buttons to indicate the presence or absence of the target within the array. The task required an initial fixation (until the spacebar press) on a fixation at the screen center followed by a brief presentation of the target for that trial (1000 ms), followed by a presentation of the search array (7000 ms). B) Search time distribution for TP trials (left, depicted in a purple gradient color) and TA trials (left, represented in a blue gradient color) across various set sizes. C) Search time (in ms, on the left) and accuracy (in %, on the right) as functions of set size for TP trials (depicted in purple) and TA trials (represented in blue). D) Search time (in ms, at the top) and accuracy (in %, at the bottom) for TP trials (illustrated in purple) and TA trials (depicted in blue) across collapsed set sizes. E) Schematic of the drift-diffusion model (DDM). F) Drift rate (v) as functions of set-size for TP trials (purple) and house trials (blue). The symbols *, **, and *** indicate statistical significance levels at p < 0.05, p < 0.01, and p < 0.001, respectively. The symbols =, /, and x represent the main effects of search efficiency, set size, and their interaction, respectively.

## Drift diffusion Model

According to the theory of memory retrieval ^17^, decision-making entails a simultaneous comparison of evidence favoring two options over time. Both the reaction time (RT) and accuracy of decisions, which are retrieved from memory, depend on the analysis of traces. Evidence accumulates gradually over time, with one aspect of the decision-making process focusing on option selection and another determining the decision endpoint. In the drift-diffusion model, evidence is incrementally collected and directed toward one of the two parallel boundaries representing the options. A decision is assumed to be reached when the difference in evidence reaches the threshold (Fig 1E) ^18^. Diffusion model analysis is based on the multi-dimensional search for an optimal set of estimates for all free parameters. Maximum Likelihood (ML) algorithms are highly efficient and are broadly applied to optimization problems for different models. In the case of diffusion models, the natural logarithms of density values (g) that calculated from predicted RT-distributions are summed over all trials *i* (with response time *RT_i_* and response *K_i_*):

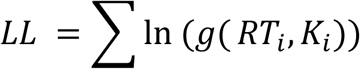

a minimum value for density of *g* = 10^−6^ is used ^19^.

In this study, the Threshold Separation (a) was equal to 1, and for Relative Starting Point (zr) 0.5 was utilized with Fast-dm software (URL link: https://www.psychologie.uni-heidelberg.de/projekt/fast-dm/) ^19^. This software enables the estimation of parameters for the drift-diffusion model by utilizing participants’ performance results and RT. The threshold of decision, drift rate, and time-related to sensory and motor processes were defined as free parameters ^18^. The rate of accumulation of information is called the drift rate (v), and it is determined by the quality of the information extracted from the stimulus. In an experiment, the value of drift rate, v, would be different for each stimulus condition that differed in difficulty ^20,21^. A higher drift rate produces faster and more accurate responses, while a lower drift rate produces slower and less accurate responses.

### 2.3 Procedure

The experiment was presented using Inquisit (version 6.6.1) on a 19-inch LCD monitor. Participants were seated approximately 60cm from the monitor. In this experiment, each trial was initiated with the display of a central cross on the screen. Subjects were explicitly instructed to fixate on the dot and press a button to signal their readiness to proceed. In trials where the target was present 50% of the time, subjects communicated their decision regarding the presence or absence of the target within the display by manually pressing a button, subsequently terminating the display. To familiarize themselves with the task and receive performance feedback, each subject underwent approximately 24 training trials. Notably, feedback on trial performance was not provided during the testing phase. The task also included an initial fixation cross, sustained until the participant pressed the spacebar, on a centrally located fixation cross on the screen. Following this fixation period, a brief presentation of the target for the current trial, lasting 1000 milliseconds, ensued. Subsequently, the search array was presented, remaining on the screen for 7000 milliseconds. To fulfill the task requirements, each participant completed around 240 trials. Participants were instructed to respond as quickly as possible without making errors. The entire task spanned 25 minutes. This study commenced on 01/04/2023 and concluded on 01/06/2024.

## Statistical tests and significance levels

A one-way ANOVA test was conducted to determine the significance of trial type, set size, age, and target location on search time, accuracy, and drift rate (v). RT, accuracy, and drift rate (v) were compared using t-tests for TP and TA trials. The significance level for all statistical analyses in this study was set at P < 0.05.

## Results

In our investigation of subjects’ performance in a conjunction visual search task using behavioral and modeling evidence. The task required subjects to identify a target among distractors following a cue presentation (target-present trials) or to signal the absence of a target if none was present (target-absent trials), as detailed in Fig. 1A and the Methods section. Our comprehensive analysis provides insights into the cognitive processes underlying visual search tasks.

### Prolonged search time and set size dependency are observed in target-absent trials compared to target-present trials in conjunction visual search

#### 1. Behavioral evidence

The analysis of search times in the current study revealed distinct patterns between target-present and target-absent trials. Participants’ responses were measured across a range of conditions, providing insights into the temporal dynamics of their cognitive processes. In target-present trials, where participants were tasked with identifying a specific target stimulus, the distribution of search times exhibited a characteristic pattern. The majority of participants demonstrated relatively faster search times, suggesting efficient processing when the target was present (Fig 1B, Supplementary Fig 1B). The central tendency of search times in these trials was indicative of the participant’s ability to quickly and accurately identify the target stimulus among distractors. Conversely, in target-absent trials, where participants had to determine the absence of the target stimulus, the distribution of search times presented a distinct profile. Search times in these trials were generally longer compared to target-present trials, reflecting the additional cognitive load associated with the decision-making process when the target was not present. Participants seemed to engage in more thorough processing, carefully scanning the presented stimuli to confirm the absence of the target (Fig 1B). The distribution of search times in both target-present and target-absent trials provides valuable insights into the cognitive mechanisms involved in visual search tasks. The observed patterns shed light on the interplay between perceptual processes, decision-making strategies, and the inherent challenges posed by the presence or absence of the target stimulus.

The insights derived from the data presented in Figure 1C provide a nuanced understanding of the intricate patterns observed in conjunction visual search. When the target was absent, participants exhibited a prolonged search time compared to trials where the target was present (t=39.04, p=.001, Fig. 1D). This discrepancy implies that the absence of a target necessitated more extensive cognitive processing, leading to an extended search duration. Conversely, in trials with a present target, participants efficiently identified its presence amidst distractors, resulting in a shorter search time. Furthermore, the examination of search time dynamics in target-absent trials revealed a noteworthy correlation with set size. As set size increased, the subjects’ search time demonstrated a consistent upward trend, indicating a heightened reliance on set size when the target was not present (main effects: f_1,7763_=2519.50, p=.001, set size: f_3,7763_=1482.27, p=.001, interaction: f_3,7763_=135.35, p=.001, Fig. 1C). However, this dependence on set size was notably diminished in target-present trials during conjunction search, suggesting a more streamlined cognitive process when the target was identified.

Moving beyond search time, our analysis delved into accuracy, uncovering compelling patterns. Participants exhibited heightened accuracy in target-absent trials, suggesting a robust ability to correctly identify the absence of the target (t=14.29, p=.001, Fig. 1D). Intriguingly, this accuracy did not exhibit a significant improvement with larger set sizes, indicating that participants maintained consistent accuracy levels regardless of the number of distractors. In stark contrast, target-present trials unveiled a more complex relationship between accuracy and set size. As set sizes increased in conjunction search, participants demonstrated a diminishing accuracy, implying that the cognitive load intensified with a greater number of distractors (main effects: f_1,8669_=212.47, p=.001, set size: f_3, 8669_=44.39, p=.001, interaction: f_3,8669_=25.04, p=.001, Fig. 1C and Supplementary Fig. 1C). This finding suggests that the ability to accurately pinpoint the target amid distractors in target-present trials was more susceptible to the challenges posed by an increasing set size.

Our study not only sheds light on the temporal dynamics of conjunction visual search but also unravels the intricate interplay between search time, accuracy, and set size in both target-absent and target-present trials.

#### 2. Modeling Evidence

In this study, results using the DDM for confirmed behavioral data revealed that revealed a significant disparity in drift rates between TP and TA trials across set sizes (t = -11.59, p < 0.001, Fig. 1F). TP trials exhibited significantly higher drift rates compared to TA trials (F_1,184_ = 221.263, p < 0.001, Fig. 1F), indicating more efficient information accumulation in target-present trials. Furthermore, our analysis demonstrated a significant dependence on set sizes, with a substantial decrease in drift rates as set sizes increased (F_3,184_ = 39.035, p < 0.001, Fig. 1F). This suggests that larger set sizes make the search task more challenging and reduce the efficiency of information processing. Additionally, a significant interaction effect was observed between trial type (TP and TA) and set sizes, emphasizing the intricate relationship between trial type and the influence of set sizes on drift rates (F_3,184_ = 3.814, p = 0.011, Fig. 1F). This indicates that the influence of set size on drift rates varies between TP and TA trials.

### An increase in target distance from the visual field center leads to a decrease in efficiency in finding targets in conjunction visual search

#### 1. Behavioral evidence

This study delves into the intricate relationship between target distance, set size, and search time during the task of target localization, as illustrated in Figure 2. The findings elucidate a compelling correlation wherein the extension of the distance from the central fixation cross corresponds to an increase in set sizes, resulting in a consistent elevation of search time (main effects: f_8,3715_=25.588, p=.001, set size: f_3,3715_=328.299, p=.001, interaction: f_24,3715_=1.797, p=.01, Fig. 2A). This trend accentuates the escalating difficulty faced by participants as they endeavor to precisely locate the target when it transitions from the central fixation cross to the peripheral boundaries of the visual field. The observed augmentation in search time with larger set sizes implies a heightened cognitive load, especially notable when the target is positioned at a greater distance from the fixation cross. These results underscore the intricate interplay between spatial factors and set size, intricately influencing the efficiency of target localization.

**Fig. 2.**
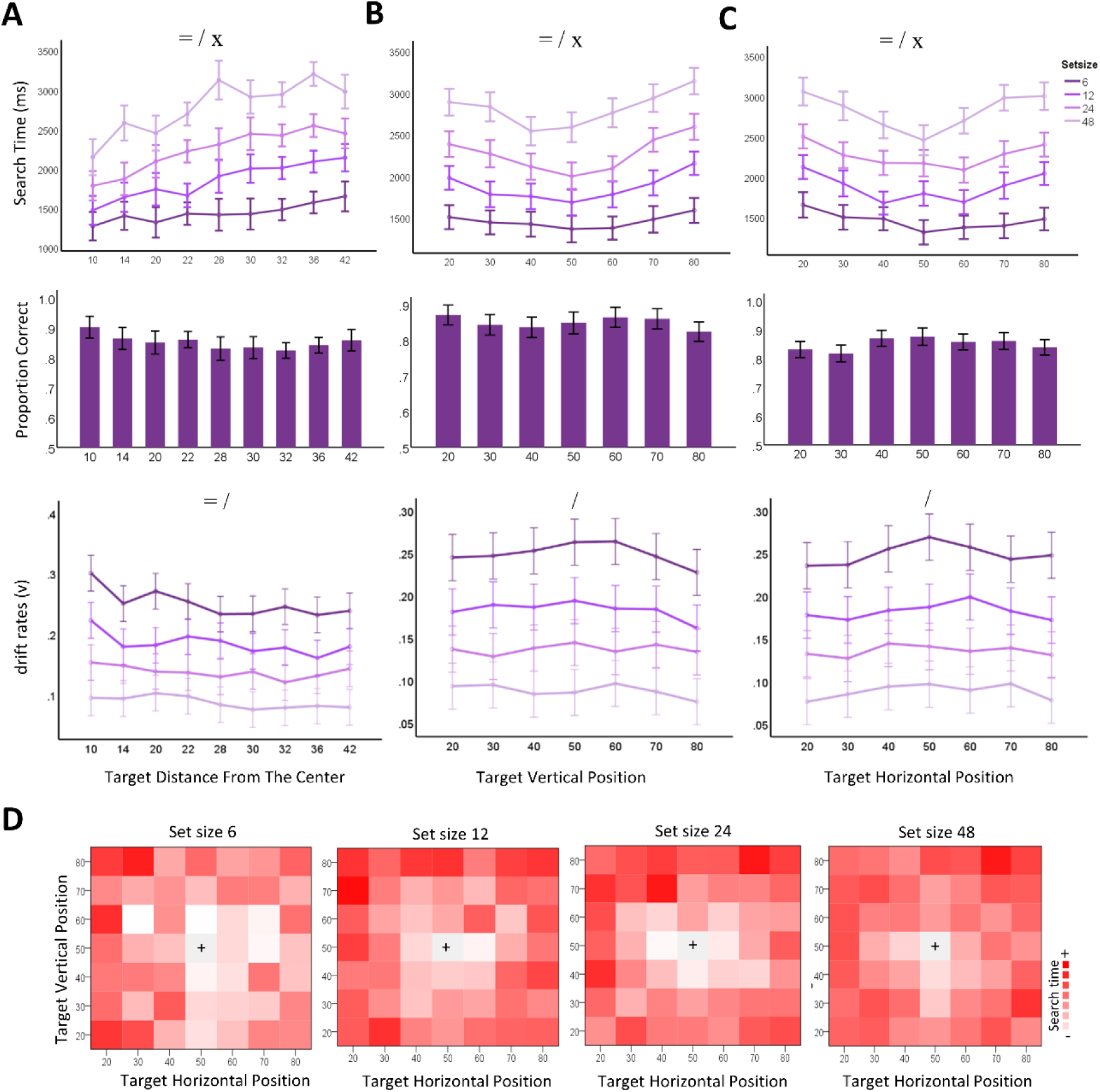
Analysis of Search Time and Accuracy in Target Detection Trials Across Varying Set Sizes and Positions. A) Search time (ms, top), accuracy (in %, middle), and drift rate (*v*, bottom) as functions of target distance from the center in TP trials across various set sizes and collapsed set sizes, respectively. B) Search time (ms, top), accuracy (in %, middle), and drift rate (v, bottom) as functions of target vertical position in TP trials across set sizes and collapsed set sizes, respectively. C) Similar format as B but for the target horizontal position. D) Heatmap illustrating the mean search time for each target position (horizontal vs. vertical) across various set sizes. Symbols *, **, and *** denote statistical significance levels at p < 0.05, p < 0.01, and p < 0.001, respectively. Symbols =, /, and x represent the main effects of search efficiency, set size, and their interaction, respectively.

Significantly, participants demonstrate reduced search times when the target is close to the center, emphasizing the inherent challenge of locating stimuli within the central visual field in both target vertical locations (main effects: f_6,3723_=17.005, p=.001, set size: f_3,3723_=368.517, p=.001, interaction: f_18,3723_=.939, p=.53, Fig. 2B, Fig. 2D, and Supplementary Fig. 2C) and in target horizontal location (main effects: f_6,3723_=14.503, p=.001, set size: f_3,3723_=367.340, p=.001, interaction: f_18,3723_=1.19, p=26, Fig. 2C, Fig. 2D). In contrast, the figure portrays an observable increase in search time for target identification as the distance from the center expands, shedding light on the dynamic nature of this correlation. In addition, the results indicated a significant difference in accuracy depending on the target distance from the center (f_8,4404_=1.946, p=.04, Fig. 2A, Supplementary Fig. 2A), as well as for the target horizontal position (p=.03, Fig. 2C, Supplementary Fig. 2B). Notably, there was no significant difference in accuracy based on the target vertical position (p=.19, Fig. 2B). Moreover, the data reveals a fascinating phenomenon—a surge in search time for target identification with an increase in distance from the fixation cross with a rise in set size. This discovery suggests that the complexity introduced by a larger set size not only impacts performance at greater distances but also influences the efficiency of target localization within the central region. Of particular note, the research brings to the forefront the intricate challenges associated with target localization, elucidating how variations in both distances from the center and set size intricately contribute to the modulation of search times.

#### 2. Modeling evidence

The DDM results for confirmed behavioral data revealed that as the target distance from the center increased, drift rates (v) decreased (F_8,108_ = 3.03, p = 0.004, Fig. 2A). This documented inverse relationship suggests that targets positioned further from the center are processed less efficiently, indicating heightened challenges in detecting and accumulating information from stimuli in the peripheral visual field. Additionally, a set size dependency on drift rates (v) was observed; as set sizes increased, drift rates (v) decreased (F_3,108_ = 197.16, p < 0.001, Fig. 2A). This significant finding underscores that larger set sizes lead to lower drift rates, illustrating that as the number of items in the visual field expands, the task’s complexity rises, resulting in diminished efficiency in processing information. Moreover, no significant interaction emerged between set size and target distance from the center, underscoring that the impact of set size on drift rates remains consistent irrespective of the target’s distance from the center (F_24,108_ = 0.514, p = 0.96, Fig. 2A). This suggests that set size and target distance independently influence drift rates, portraying their distinct roles in shaping processing efficiency.

Furthermore, our DDM results showcased heightened drift rates (v) when the target was near the center, emphasizing the inherent challenge of locating stimuli within the central visual field. This effect persisted across both target vertical and horizontal locations (main effects: F_6,84_ = 1.108, p = 0.36; set size: F3,84 = 176.88, p < 0.001; interaction: F_18,84_ = 0.26, p = 0.99, Fig. 2B, Fig. 2C), indicating superior processing efficiency for targets positioned closer to the center. This enhanced efficiency may stem from heightened visual acuity and intensified attention within the central visual field.

### Age-related dynamics negatively affect efficiency in conjunction visual search

#### 1. Behavioral evidence

The findings of the study indicate a noteworthy association between age and performance in conjunction search tasks. As subjects’ age increases, there is a discernible escalation in the search time required to reject the presence of the target in trials where the target is absent (main effects: f_1,7597_=1507.891, p=.001, age: f_4,7597_=25.53, p=.001, interaction: f_4,7597_=2.546, p=.03, Fig. 3A and Supplementary Fig. 3A). This is particularly evident when contrasting it with the relatively more efficient task of finding the target in trials where it is present. The elevated search time in the absence of the target suggests a growing difficulty among subjects in declaring its non-existence as they age. The age-related effect remains consistent across various set sizes in both target-present trials (main effect: f_4,3654_=32.145, p=.001; set size: f_3,3654_=350.002, p=.001; interaction: f_12,3654_=1.068, p=.38, Fig. 3B) and target-absent trials (main effect: f_4,3654_=29.450, p=.001; set size: f_3,3654_=1246.384, p=.001; interaction: f_12,3654_=3.81, p=.001, Fig. 3C). While a significant interaction between age and set size was observed in target-absent trials, no significant interaction was found between age and set size in target-present trials. The dependence on set size is more prominent in absent trials compared to trials where the target is present, further emphasizing the challenges that arise with age in conjunction search tasks (Fig. 3, Supplementary Fig. 3A-B).

**Fig. 3.**
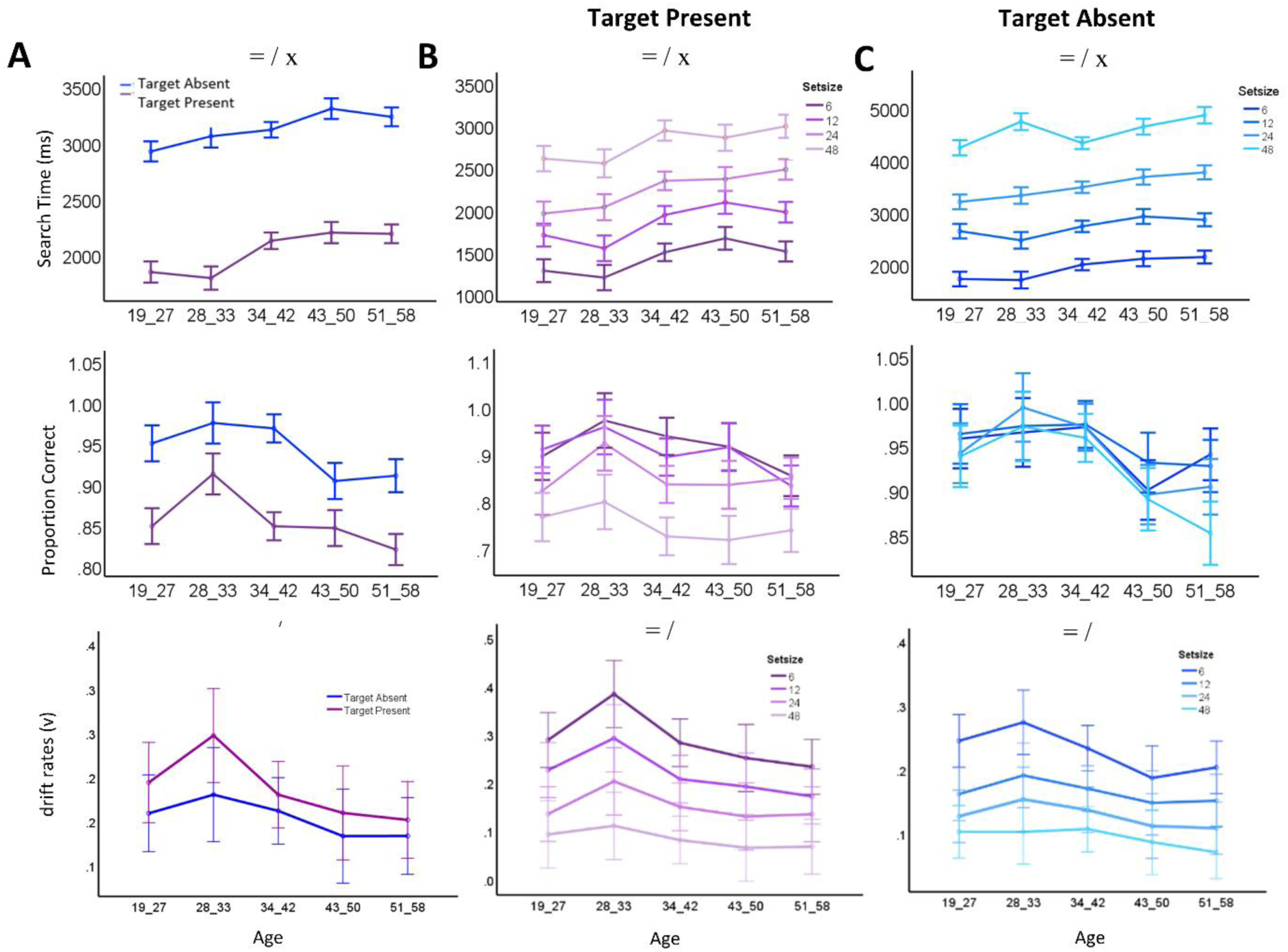
Age-Related Analysis of Search Time and Accuracy in Target Detection Trials Across Set Sizes and Trial Types (TP and TA). A) Search time (ms, top) and accuracy (%, middle), and drift rate (v, bottom) as functions of age in TP (purple) and TA (blue) trials across collapsed set sizes. B) Search time (ms, top) and accuracy (%, middle), and drift rate (v, bottom) as functions of age in TP trials across set sizes. C) Similar format as B but for the TA trials. Symbols =, /, and x represent the main effects of search efficiency, set size, and their interaction, respectively.

Moreover, the study delves into the accuracy of subjects in both trials. It is observed that, with increasing age, subjects exhibit greater accuracy in rejecting the presence of the target in trials where it is absent, in contrast to their performance in locating the target in trials where it is present. This suggests that, as subjects age, their ability to accurately declare the absence of a target improves (main effects: f_1,8480_=163.279, p=,001, age: f_4,7597_=15.01, p=.001, interaction: f_4,7597_=3.305, p=.01, Fig. 3A and Supplementary Fig. 3B). Interestingly, the analysis extends to the impact of set sizes on accuracy. While the accuracy in declaring the absence of the target remains higher with age, there is a notable decrease in accuracy as set sizes increase (main effect: f_4,4152_=17.58, p=.001, set size: f_3, 4152_=3.23, p=.02, interaction: f_12, 4152_=1.120, p=.339, Fig. 3B). This decline is evident in both absent and target-present trials (main effect: f_4,3654_=6.75, p=.001, set size: f_3,3654_=46.415, p=.001, interaction: f_12,3654_=1.222, p=.26, Fig. 3C) in conjunction search task, highlighting the nuanced interplay between age, set size, and task performance.

#### 2. Modeling evidence

For confirmed behavioral results, DDM results showed that there was a significant difference observed between drift rate groups in TP and TA trials at different ages, reflecting that in target-absent trials, the drift rates were higher than in target-present trials (F_1,101_ = 4.87, p = 0.02, Fig. 3A). This suggests that individuals tend higher drift rates when the target is absent, potentially indicating a greater cognitive effort in processing the absence of a target compared to its presence. However, the drift rates demonstrated a significant dependence on age (F_4,101_ = 2.535, p = 0.04, Fig. 3A), suggesting that with increased age, drift rates decreased. This indicates a potential age-related refinement in cognitive processing efficiency during visual search tasks. Additionally, no significant interaction effect was observed between the trial type (TP and TA) and age, emphasizing the intricate relationship between the trial type and the influence of age on the drift rate (F_4,101_ = 0.34, p = 0.86, Fig. 3A). In addition to this, our DDM results demonstrate decreased drift rates (v) as age increased in both target-present trials (main effects: F_4,35_ = 5.24, p = 0.002, set size: F_3,35_ = 41.22, p < 0.001, interaction: F_12,35_ = 0.37, p = 0.96, Fig. 3B) and target-absent trials (main effects: F_4,35_ = 3.09, p = 0.02, set size: F3,35 = 35.11, p < 0.001, interaction: F_12,35_ = 0.26, p = 0.99, Fig. 3C). This suggests that with increasing age, individuals exhibit more refined cognitive processes, resulting in decreased drift rates during visual search tasks. The observed patterns in both trial types imply a consistent age-related improvement in processing efficiency, potentially reflecting cognitive maturation or adaptation to task demands over time.

## Discussion

This study investigates subjects’ performance in a conjunction visual search task via behavioral and modeling evidence. Participants were tasked with identifying a target amidst distractors (target-present trials) or signaling its absence (target-absent trials). The analysis revealed distinct patterns in search times, with faster and more efficient processing in target-present trials and prolonged search times in target-absent trials, indicating additional cognitive load (Fig. 1). Examination of search time dynamics uncovered correlations with set size, emphasizing heightened reliance in target-absent trials. Accuracy metrics revealed consistent accuracy in target-absent trials, contrasting with diminishing accuracy in target-present trials with increasing set sizes (Fig. 1). The study additionally explored the impact of target distance on search time, emphasizing challenges in target localization, with increased difficulty observed at greater distances from the central visual field (Fig. 2). Age-related dynamics demonstrated an escalation in search time with age, particularly in target-absent trials, emphasizing challenges in declaring non-existence. Despite age-related difficulties, accuracy in declaring absence improved with age, showcasing a nuanced interplay between age, set size, and task performance (Fig. 3).

Our results offer valuable insights into the complexities of conjunction visual search, shedding light on the nuanced patterns observed in participants’ behavior. These findings are in line with previous research that has explored the intricacies of visual search processes. For instance, Wolfe and Horowitz ^22^ have emphasized the significance of understanding search times and accuracy metrics in various search tasks, highlighting the role of cognitive processes in target detection. In target-absent trials, participants demonstrated a prolonged search time, aligning with the idea that the absence of a target requires more extensive cognitive processing. This observation is consistent with the notion that searching for an absent target involves exhaustive scanning of the visual field ^23^. The correlation between search time and set size in target-absent trials is reminiscent of the set size effect, a phenomenon widely discussed in the literature ^10^. As set size increased, participants’ search time rose, indicating a heightened reliance on set size when the target was not present. Contrastingly, in target-present trials during conjunction search, participants exhibited efficient target identification with shorter search times. This finding resonates with the notion of parallel processing during feature conjunction searches, where the presence of a unique feature facilitates quicker target identification ^24^. The reduced dependence on set size in target-present trials suggests a more streamlined cognitive process, highlighting the role of feature integration in facilitating efficient visual search ^10^. The examination of accuracy metrics further contributes to the nuanced understanding of participants’ performance. In target-absent trials, heightened accuracy irrespective of set size indicates a robust ability to correctly identify the absence of the target. This aligns with previous studies emphasizing the importance of efficient rejection of distractors in visual search tasks ^25^. However, the lack of improvement in accuracy with larger set sizes in target-absent trials suggests that participants maintained a consistent ability to reject distractors regardless of their number. In target-present trials, the diminishing accuracy with increasing set sizes reflects the challenges posed by a greater number of distractors during conjunction search. This finding is in line with the idea that the integration of features becomes more error-prone as the complexity of the visual scene increases^24^. The intricate interplay between accuracy and set size in target-present scenarios highlights the vulnerability of accurate target identification when faced with an elevated cognitive load.

The presented study delves into the complex relationship between target distance, set size, and search time, shedding light on the intricate dynamics of target localization. The observed correlation, illustrated in Figure 2, reveals a noteworthy pattern where an increase in target distance from the central fixation cross corresponds to larger set sizes, resulting in a consistent elevation of search time across all experimental conditions. These findings suggest that as the target transitions from the central fixation cross to the peripheral boundaries of the visual field, participants face an escalating difficulty in precisely locating the target, manifesting as increased search times. This observed augmentation in search time with larger set sizes implies an elevated cognitive load, particularly pronounced when the target is positioned at a greater distance from the fixation cross. The results underscore the intricate interplay between spatial factors and set size, intricately influencing the efficiency of target localization. This is consistent with previous research that has highlighted the role of cognitive load in visual search tasks ^23^. Interestingly, the study also reveals reduced search times when the target is in close proximity to the center, emphasizing the inherent challenge of locating stimuli within the central visual field (Fig. 2B-D). This finding aligns with prior research suggesting that the central visual field is processed more efficiently than the peripheral regions ^26^. In contrast, an observable increase in search time for target identification is noted as the distance from the center expands, highlighting the dynamic nature of the correlation between target distance and search time. Furthermore, the data illuminates a fascinating phenomenon—a surge in search time for target identification with an increase in distance from the fixation cross coupled with a rise in set size. This discovery suggests that the complexity introduced by a larger set size not only impacts performance at greater distances but also influences the efficiency of target localization within the central region. These results align with studies emphasizing the interactive effects of set size and spatial factors on visual search performance ^10^.

The study under consideration sheds light on the intricate relationship between age and performance in conjunction search tasks, revealing compelling insights into the impact of aging on various aspects of task execution. The most prominent finding revolves around the discernible association between age and search time, particularly in trials where the target is absent. The observed escalation in search time with increasing age suggests a growing difficulty among subjects in promptly rejecting the presence of the target when it is not there. This finding aligns with existing literature on cognitive aging, where processing speed and efficiency tend to decline with age ^27^. The study further highlights that this age-related effect is not confined to specific set sizes but remains consistent across various set sizes. This is a noteworthy observation, emphasizing that the challenges associated with aging in conjunction search tasks persist regardless of the complexity of the visual field. The dependence on set size, as revealed by the study, is more pronounced in trials where the target is absent compared to trials where the target is present. This underscores the challenges individuals face in efficiently processing and discriminating non-targets as they age, especially in scenarios requiring the rejection of potential distractors. Similar findings have been reported in studies examining visual search tasks and aging ^13^, supporting the notion that age-related changes in attentional processes contribute to the observed difficulties. Moving beyond search time, the study delves into the accuracy of subjects in both scenarios. The intriguing observation is that, as subjects age, they exhibit greater accuracy in rejecting the presence of the target in absent trials compared to locating the target in present trials. This counterintuitive pattern suggests a potential compensatory mechanism, wherein older individuals may develop enhanced skills in accurately declaring the absence of a target as a way to cope with the challenges posed by aging ^28^. The examination of accuracy in relation to set sizes adds another layer of complexity to the findings. While accuracy in declaring the absence of the target remains higher with age, there is a notable decrease in accuracy as set sizes increase. This nuanced interaction underscores the intricate interplay between age, set size, and task performance. The decline in accuracy with larger set sizes implies that the challenges associated with visual discrimination and attentional selection become more pronounced in complex visual environments, even for older individuals who demonstrate improved accuracy in rejecting the target’s presence.

In summary, this study provides comprehensive insights into the complexities of conjunction visual search, unraveling distinct patterns in participants’ behavior across various conditions. Prolonged search times in target-absent trials, influenced by set size, highlight the cognitive load associated with exhaustive scanning, while efficient target identification in target-present trials underscores streamlined cognitive processes. The examination of target distance, set size, and search time reveals the intricate dynamics of target localization, emphasizing the central visual field’s efficiency and the impact of spatial factors. Additionally, age-related increases in search time, coupled with nuanced accuracy patterns, contribute valuable insights into the interplay between aging, set size, and task performance in conjunction visual search, ultimately enriching our understanding of this cognitive process. Finally, the ability to detect the presence or absence of a target holds paramount importance in decision-making, particularly when our perceptions are susceptible to influence from belief systems ^29,30^. Therefore, it is advisable to employ visual search as a cognitive function examination to scrutinize many of these beliefs, including paranormal beliefs, which may be afflicted by cognitive defects ^29–38^.

## Acknowledgments

We thank participants for their helpful cooperation.

## Ethical consideration

This study is supported by the *School of Cognitive Science at the Institute for Research in Fundamental Sciences (IPM)*. Our research strictly followed the ethical guidelines outlined in the Declaration of Helsinki. Before participating in the study, all participants provided written informed consent.

## Data and materials availability

All relevant data are within the paper and its Supporting information files.

---

**Supplementary Fig 1.**
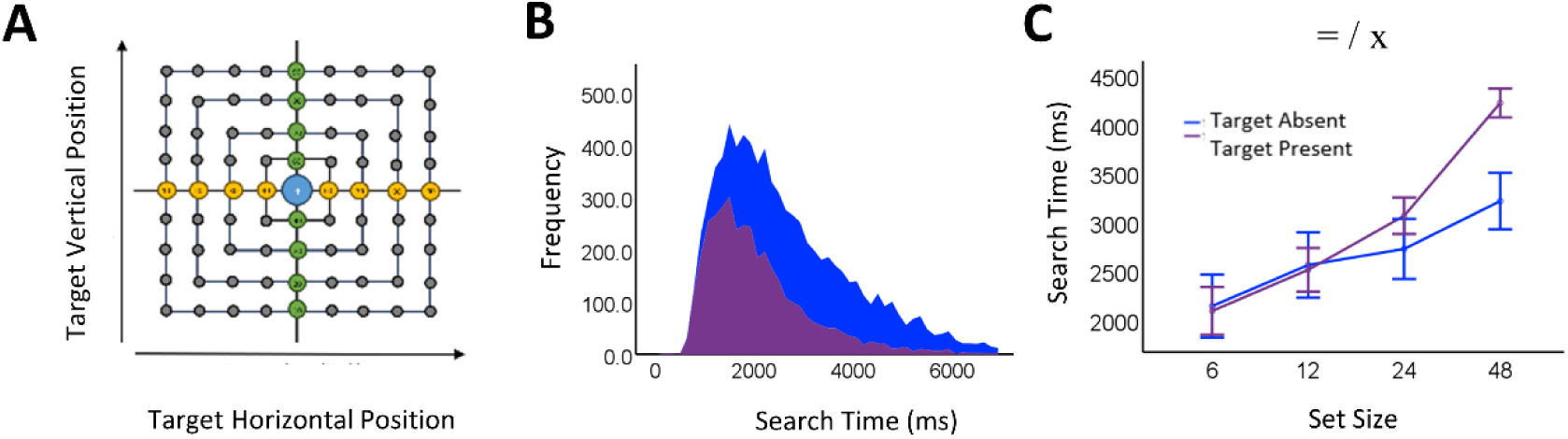
A) Target position (horizontal vs. vertical) in the visual search task during TP trials. B) Search time (ms) distribution for TP trials (in purple) and TA trials (in blue) across collapsed set sizes. C) Search time (ms, on the right, main effects: f_1,898_=11.081, target present: f_3,898_=58.282, p=.001, interaction: f_3,898_=7.986, p=.001) as a function of set size for TP trials (depicted in purple) and TA trials (represented in blue) during incorrect trials.

**Supplementary Fig 2.**
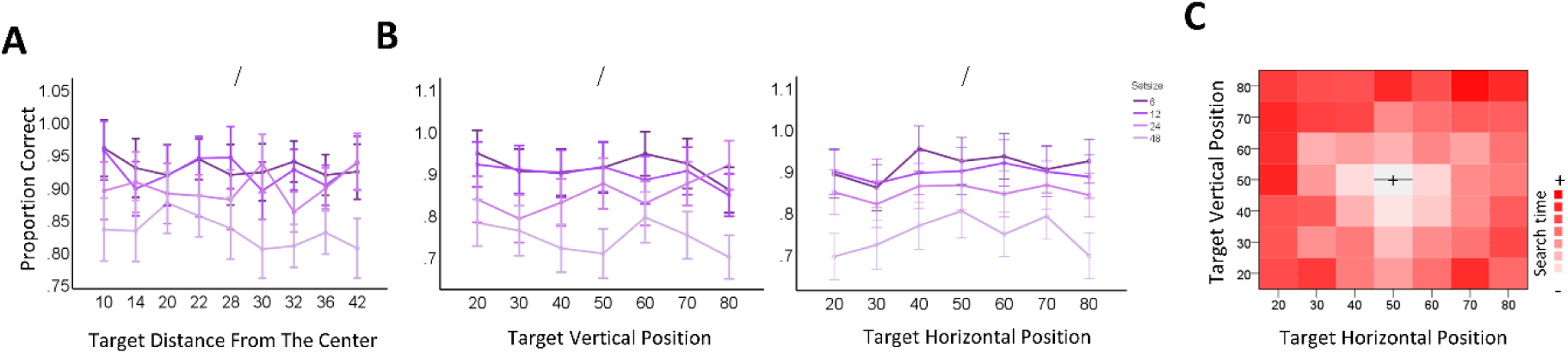
A) Accuracy (%, on the left, main effects: f_8,8641_=1.002, p=.43, set size: f_3,8641_=41.795, p=.001, interaction: f_24,8641_=1.220, p=.21) as a function of target distance from the center in TP trials across various set sizes. B) Accuracy in vertical position in TP trials across various set sizes (%, in the left, main effects: f_6,4385_=1.215, p=.29, set size: f_3,4385_=49.807, p=.001, interaction: f_18,4395_=1.522, p=.07) as a function of target, and accuracy in the target horizontal position (%, on the right, main effects: f_6,4385_=2.271, p=.03, set size: f_3,4385_=48.762, p=.001, interaction: f_18,4385_=.650, p=.86). C) Heatmap illustrating the mean search time for each target position (horizontal vs. vertical) across collapsed set sizes. Symbols =, /, and x represent the main effects of search efficiency, set size, and their interaction, respectively.

**Supplementary Fig 3.**
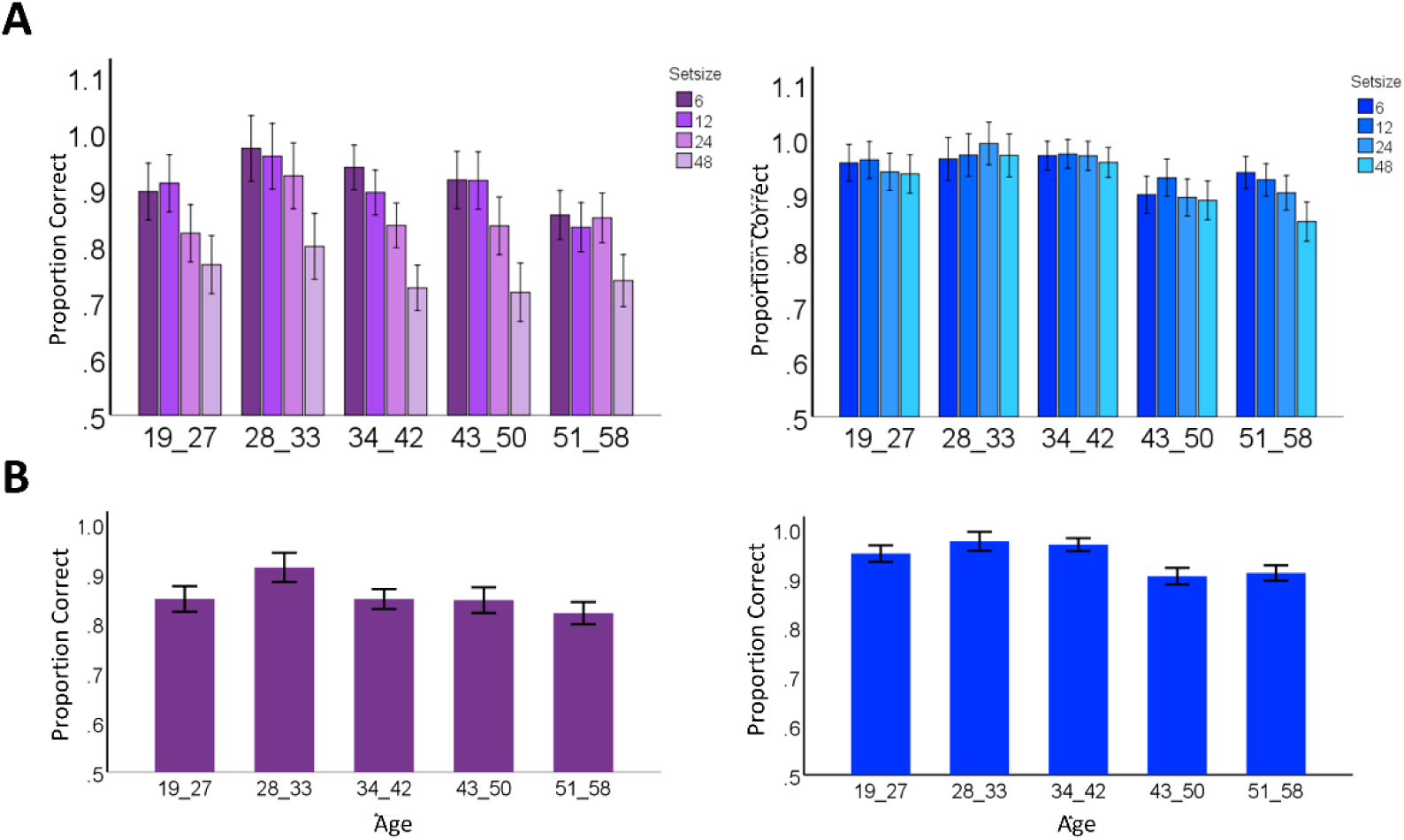
A) Accuracy (%, top, main effects: f_4,4298_=6.758, p=.001, set size: f_3,4298_=46.415, p=.001, interaction: f_12,4298_=1.222, p=.001) as a function of age in TP trials (left) and Accuracy (%, on the top, main effects: f_4,4152_=17.587, p=.001, set size: f_3,4152_=3.235, p=.021, interaction: f_12,4152_=1.120, p=.33) TA trials (right) across various set sizes. B) Accuracy (%, bottom, main effects: f_4,4313_=6.244, p=.001) as a function of age in TP trials (left) and (%, bottom, main effects: f_4,4167_=16.282, p=.001) TA trials (right) across collapsed set sizes.

